# Developmental Variations in Recurrent Spatiotemporal Brain Propagations from Childhood to Adulthood

**DOI:** 10.1101/2025.02.04.635765

**Authors:** Kyoungseob Byeon, Hyunjin Park, Shinwon Park, Jon Cluce, Kahini Mehta, Matthew Cieslak, Zaixu Cui, Seok-Jun Hong, Catie Chang, Jonathan Smallwood, Theodore D. Satterthwaite, Michael P. Milham, Ting Xu

**Affiliations:** Child Mind Institute, New York, NY, United States; School of Electronic and Electrical Engineering, Sungkyunkwan University, Suwon, South Korea; IBS Center for Neuroscience Imaging Research, Sungkyunkwan University, Suwon, South Korea; Penn Lifespan Informatics and Neuroimaging Center (PennLINC), Perelman School of Medicine, University of Pennsylvania, Philadelphia, PA, United States; Penn-CHOP Lifespan Brain Institute, Children’s Hospital of Philadelphia and Perelman School of Medicine, University of Pennsylvania, Philadelphia, PA, USA; Beijing Institute for Brain Research, Chinese Academy of Medical Sciences & Peking Union Medical College, Beijing, China; Chinese Institute for Brain Research, Beijing, China; Department of Biomedical Engineering, Sungkyunkwan University, Suwon, South Korea; Department of Intelligent Precision Healthcare Convergence, Sungkyunkwan University, Suwon, South Korea; Departments of Electrical and Computer Engineering, Computer Science, and Biomedical Engineering, Vanderbilt University, Nashville, TN, United States; Department of Psychology, Queens University, Kingston, ON, Canada; Nathan S. Kline Institute for Psychiatric Research, Orangeburg, NY, United States

**Keywords:** Neurodevelopment, fMRI, cortical development, dynamic brain activity, adolescence, top-down processing

## Abstract

The brain undergoes profound structural and functional transformations from childhood to adolescence. Convergent evidence suggests that neurodevelopment proceeds in a hierarchical manner, characterized by heterogeneous maturation patterns across brain regions and networks. However, the maturation of the intrinsic spatiotemporal propagations of brain activity remains largely unexplored. This study aims to bridge this gap by delineating spatiotemporal propagations from childhood to early adulthood. By leveraging a recently developed approach that captures time-lag dynamic propagations, we characterized intrinsic dynamic propagations along three axes: sensory-association (S-A), ‘task-positive’ to default networks (TP-D), and somatomotor-visual (SM-V) networks, which progress towards adult-like brain dynamics from childhood to early adulthood. Importantly, we demonstrated that as participants mature, there is a prolonged occurrence of the S-A and TP-D propagation states, indicating that they spend more time in these states. Conversely, the prevalence of SM-V propagation states declines during development. Notably, top-down propagations along the S-A axis exhibited an age-dependent increase in occurrence, serving as a superior predictor of cognitive scores compared to bottom-up S-A propagation. These findings were replicated across two independent cohorts (N = 677 in total), emphasizing the robustness and generalizability of these findings. Our results provide new insights into the emergence of adult-like functional dynamics during youth and their role in supporting cognition.

## Introduction

The brain is a dynamic system with continuously evolving neural activity that interacts across networks. Within the dynamic landscape, temporal synchronization between brain networks exhibits rich time-varying features, involving rapid reconfigurations into transient whole-brain states and forming spatiotemporal patterns over time (Aquino et al., 2012; Calhoun et al., 2014; Chang & Glover, 2010; Griffa et al., 2023; Vidaurre et al., 2017). Accumulating evidence describes recurrent spatiotemporal patterns, detectable using a range of recording and imaging techniques in humans, nonhuman primates, and rodents (Griffa et al., 2023; Y. Gu et al., 2021; Majeed et al., 2011; Meyer-Baese et al., 2022). These dynamic changes may index ongoing cognition and arousal, and reflect the brain’s continuous adaptation to internal and external environment – which is critical for maintaining cognitive flexibility (Bolt et al., 2022; Gutierrez-Barragan et al., 2022; Raut et al., 2021). Importantly, recent work has identified these dynamic fluctuations as a combination of instantaneous standing waves and time-lagged traveling waves propagating across functionally defined networks (Bolt et al., 2022). In adults, the topography of these waves prominently features spatiotemporal propagations along sensorimotor-association (S-A) and ‘task-positive’-’task-negative’ axes (Bolt et al., 2022). However, it remains largely unknown how these spatiotemporal patterns emerge from childhood to early adulthood and contribute to cognitive development.

During childhood to adolescence, the brain undergoes profound structural and functional changes that shape cognition, emotion, and behavior (Tau & Peterson, 2010). Substantial evidence suggests that the brain development during this period is heterochronic across diverse brain regions. Early maturation primarily occurs in unimodal areas, including sensorimotor, visual and auditory cortices, succeeded by the higher-order association cortices that support advanced cognitive functions such as semantic processing, decision-making, problem-solving, and socioemotional cognition (Baum et al., 2022; Gao et al., 2020; Ito et al., 2020; Murray et al., 2014; Sydnor et al., 2021, 2023). Concurrently, functional brain organization also exhibits heterochronic changes during this period. In childhood, functional connectivity is predominantly organized among different unimodal regions, likely to facilitate sensory and motor experiences during the early stages of life. By adolescence, however, there is a notable shift towards an increased prominence of the functional organization along the S-A axis (Dong et al., 2021; Xia et al., 2022). Despite these advances, studies investigating functional dynamics in youth have largely relied on instantaneous synchronization with zero-lag approaches, which may not capture the whole-brain traveling waves and their propagations across networks (Bolt et al., 2022). As a result, how the dynamic propagations emerge and shape the functional organization during childhood to early adulthood remains poorly understood.

Here, we investigate age-related changes in brain dynamic propagations during childhood and early adulthood. To capture the time-varying nature of propagation across the developmental samples previously unattainable, we leveraged Complex Principal Component Analysis (CPCA), a recently developed method designed to decompose dynamic systems as a mixture of traveling and standing waves (Bolt et al., 2022). CPCA is capable of identifying zero-lag and time-lag inter-regional synchrony by detecting recurring spatiotemporal propagations that are representative of prior studies on temporal dynamics (e.g. quasi-periodic patterns) (Bolt et al., 2022; Majeed et al., 2011; Thompson et al., 2014; Yousefi & Keilholz, 2021). Prior work using this method identified oscillating waves in young adults spanning sensorimotor to association networks (S-A) (Bolt et al., 2022). Beyond this, they also captured an additional axis of propagation between task-positive and default networks (TP-D) (Bolt et al., 2022). Building on these findings, we characterized these propagations and investigated their maturation from childhood to early adulthood. We hypothesized that these spatiotemporal propagations develop progressively from childhood to early adulthood, maturing to resemble adult-like brain dynamics. We recognized that these time-resolved propagations share similar spatial patterns with the cortical gradients of static functional connectivity. To examine this relationship, we further compared the time participants spent in each propagation state with the explanatory power of corresponding functional gradients to reveal how dynamic propagations contribute to the development of static functional gradients during this period. Next, considering that the direction of propagations (e.g. top-down versus bottom-up) might vary between youth and adults (Pines et al., 2023), we examined age-related variations in the prevalence of specific propagation direction during this period. We demonstrated that these propagations can predict individual differences in cognitive performance. Finally, we validated the robustness and generalizability of our findings, including the characterization of dynamic propagations and developmental trajectories, through replication in an independent dataset.

## Results

### Reliable recurring dynamic propagations were characterized from childhood to adulthood

We first characterized propagation components, which represent distinct spatiotemporal patterns that dynamically evolve brain-wide over time, using CPCA (Bolt et al., 2022). Unlike traditional dynamic techniques, which confines dynamic states within a fixed window size, CPCA flexibly tracks recurring dynamic cycles in the phase domain from 0 to 2*π*. Whole-brain fMRI signals are decomposed into recurring spatiotemporal patterns, propagating across different brain regions from the starting point (phase 0 radians) to the midpoint (phase *π* radians), and then completes a full circle back to the initial starting phase (phase 2*π* or 0 radians). To examine developmental changes, we separately extracted the propagation components in developmental datasets (HCP-D, N = 408, 215 females, age range: 8-21 years) and middle-aged adult reference dataset (HCP-A, N = 399, 225 females, age range: 36-64 years). The HCP-A dataset was selected as a reference due to its identical acquisition parameters to HCP-D. In both datasets, the first three propagation components explained over 50% of the total variance of resting-state fMRI dynamics. To illustrate the propagations across the cortex at different cycle stages, we visualized the spatiotemporal patterns of each component over the phase 0 to 2*π* sampled at specific phase points (Fig. 1A). The first three group propagation components in the reference dataset revealed distinct spatiotemporal dynamics. The first propagation spans the cortex between the somatosensory and transmodal association networks (Pattern S-A). The second propagation captures a cycle between the task-positive networks (i.e. frontoparietal and attention networks) and the default mode network (Pattern TP-D). The third component propagates between somatomotor and visual networks (Pattern SM-V). Each of these three spatiotemporal patterns explained over 10% of the variance for both HCP-A and HCP-D (Fig. 1B). Of note, the raw order of these components was determined by the explained variance ratio (EVR), which varies across individuals and between youth and adult cohorts. To enable inter-individual comparability of spatiotemporal patterns for both within the HCP-D cohort and relative to the adult reference sample (HCP-A), we used orthogonal alignment to project the individual-specific components onto a shared dimensional space from adult reference. At the group level, each of the spatiotemporal patterns exhibited high similarity (*r* > 0.85) across youth and adult cohorts (Fig. 1C, diagonal). At the individual level, the spatiotemporal patterns were replicable across sessions within individuals, achieving high test-retest reliability (Fig. 1D) with discriminability over 0.9 for Pattern S-A and TP-D, and over 0.85 for Pattern SM-V.

**Figure 1.**
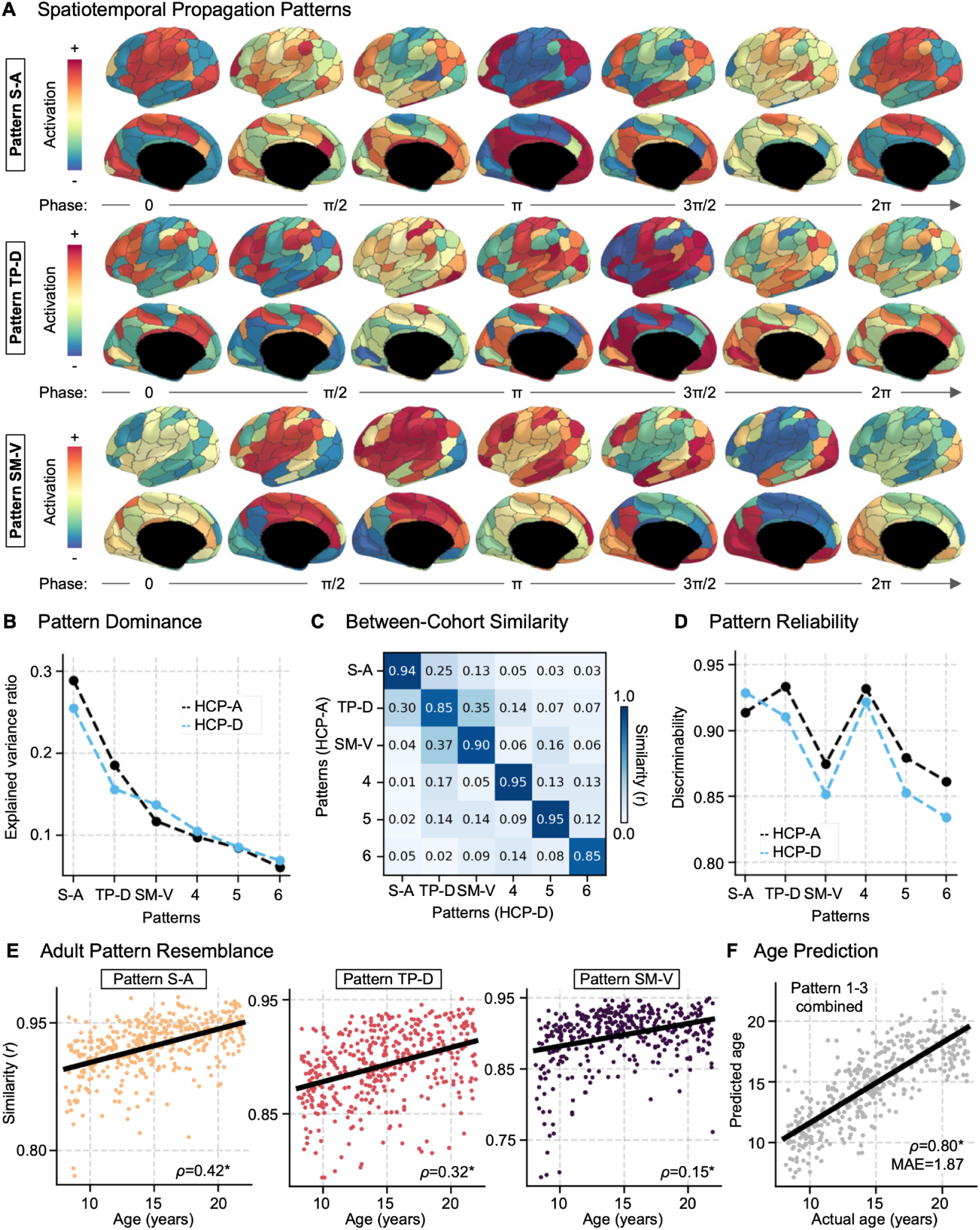
Spatiotemporal propagation patterns and its neurodevelopmental change from children to early adulthood. **(A)** The first three propagation patterns derived from the reference cohort (HCP-A), representing group-level reference propagation patterns. Each row displays a full propagation cycle for the recurring spatiotemporal patterns: sensorimotor to association (S-A), task-positive to default mode networks (TP-D), and somatomotor to visual networks (SM-V). The patterns are depicted through their temporal phase cycle, ranging from 0 to 2*π*. **(B)** Explained variance ratios of the first six propagation patterns from CPCA. The light blue line represents the youth cohort (HCP-D) and the dark line represents the reference adult cohort (HCP-A). **(C)** Between cohort similarity matrix showing the pairwise Pearson’s correlation of the propagation patterns across youth (HCP-D) and adult (HCP-A) propagation patterns. **(D)** Reliability of propagation patterns, assessed by the discriminability for HCP-D and HCP-A cohorts. **(E)** Age-related similarity of propagation patterns to adult reference. Dots represent the spatial correlations of the propagation pattern between individuals in the youth cohort and the group-level adult reference. The regression line and correlation values show how the resemblance of youth to adult patterns changes with age, with statistical significance noted (* for *pFDR* < 0.05). **(F)** Age prediction using the first three dynamic patterns. A combination of the first three dominant propagation patterns in the PLSR model predicts age with a Spearman’s correlation ρ of 0.81 and mean absolute error (MAE) of 1.87 years.

### Maturation of spatiotemporal patterns in youth towards adult-like dynamics

Having demonstrated reproducible and reliable spatiotemporal patterns in individuals in youths and adults, we examined their developmental effect by comparing individual-specific propagation patterns obtained from childhood to early adulthood cohort HCP-D with the middle-aged adult reference patterns (Fig. 1E). Overall, the propagation patterns are highly similar between youths and adults (Pattern S-A, mean *r* = 0.93 ± 0.02; Pattern TP-D: *r* = 0.89 ± 0.03; Pattern SM-V: *r* = 0.90 ± 0.04). Notably, the similarity increased significantly with age for all three patterns, controlling for sex and head motion (Pattern S-A: *ρ* = 0.42; Pattern TP-D: *ρ* = 0.32; Pattern SM-V: ρ = 0.15, all *p* < 0.05) from childhood to early adulthood, indicating a progressive refinement of functional dynamics toward an adult-like spatiotemporal pattern during this period. To further assess the extent to which these three dynamic propagations collectively explain age-related neurodevelopmental changes during this period, we employed a partial least squares regression (PLSR) model with five-fold cross-validation to estimate individual ages using three propagation patterns and achieved high accuracy (Fig. 1F, *ρ* = 0.80 and Mean Absolute Error = 1.87 years).

### Individuals spend more time in Patterns S-A and TP-D but less time in Pattern SM-V from childhood to early adulthood

Next, we examined the developmental effect on the time participants spent in each propagation state. We first defined the intrinsic moment-to-moment dominant state based on the strength of the propagation patterns. The dominant state was determined using a winner-take-all approach, selecting the pattern with the highest propagation strength (temporal magnitude) at each time point. We counted the time points of the dominant state and normalized it by the total number of time points in the scan, referring to this measure as the occurrence ratio (Fig. 2A). Time points where none of the first three patterns were dominant were categorized as the “other patterns” state. We then grouped participants into three developmental stages (children: 8-12 years, adolescents: 13-17 years, early adulthood: 18-21 years) and compared the occurrence ratio of each dynamic state across these developmental periods (Fig. 2C). This approach allows for identifying the stage-specific differences linked to developmental milestones and might capture stage-specific changes tied to key neurodevelopmental transitions. For all three propagations, the occurrence ratio showed significant group differences across developmental stages (*F_(3,803)_* = 43.84, *p* < 0.01 for Pattern S-A; *F_(3,803)_* = 35.09, *p* < 0.01 for Pattern TP-D; and *F_(3,803)_* = 35.88, *p* < 0.01 for Pattern SM-V). The occurrence ratio of Pattern S-A significantly increased across all age stages from childhood to adolescence (*t_309_* = 3.33, *p* < 0.01, effect size = 0.38), from adolescence to early adulthood (*t_267_* = 2.81, *p* < 0.01, effect size = 0.36), and from early adulthood to middle adulthood (*t_494_* = 3.04, *p* < 0.01, effect size = 0.34). Pattern TP-D exhibited no significant change of state occurrence ratio from childhood to early adulthood but a significant increase between early adulthood to middle-aged adulthood (*t_494_* = 4.71, *p* < 0.01, effect size = 0.53). In contrast, Pattern SM-V decreased its occurrence ratio starting from adolescence (adolescence to early adulthood: *t_267_* = −3.77, *p* < 0.01, effect size = −0.48, early adulthood to middle adulthood: *t_494_* = −3.25, *p* < 0.01, effect size = −0.37).

**Figure 2.**
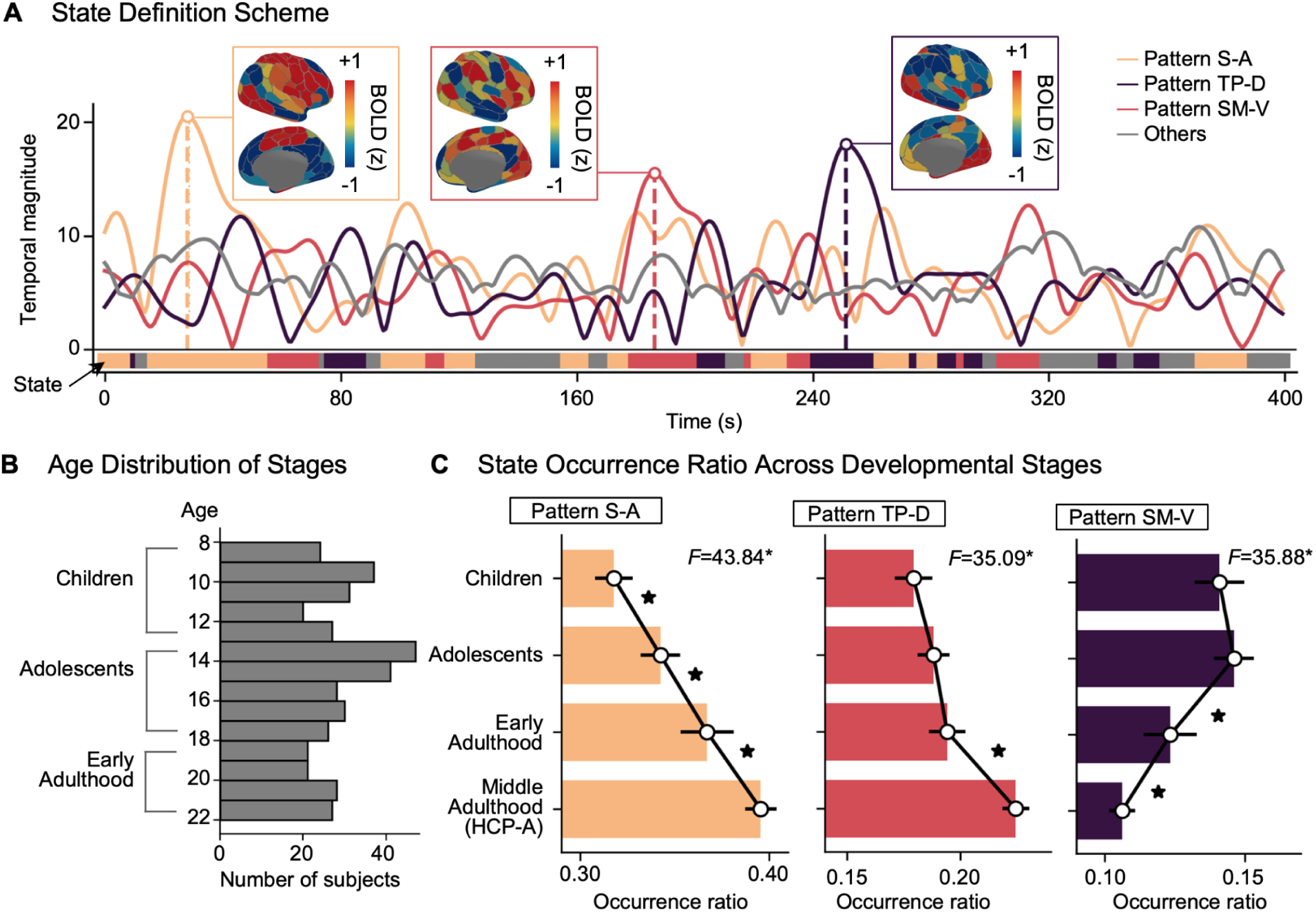
Individuals spend more time in Patterns S-A and TP-D but less time in Pattern SM-V from childhood to adulthood. **(A)** Dynamic state definition scheme. The strength of dynamic patterns at each time point from an example individual is presented. The temporal magnitude (y-axis) reflects the strength of the dynamic propagation pattern over time. At each time point, the pattern with the highest magnitude is assigned as the dominant state. **(B)** Age distribution of participants across developmental stages (children: 8-12 years, adolescents: 13-17 years, and early adulthood: 18-22 years) in the HCP-D cohort. **(C)** The proportion of time spent (i.e. occurrence ratio) is plotted for different developmental stages: children, adolescents, early adulthood in HCP-D, and middle-aged adulthood reference (HCP-A). Statistical analyses include a one-way ANOVA across the four age groups to assess overall developmental effects, followed by pairwise t-tests between consecutive age stages to examine specific transitions. The significance is denoted by asterisks (*: *p*FDR < 0.05).

### Spatiotemporal propagations reflect both dynamic and static functional organization

Notably, the spatial patterns of S-A, TP-D, and SM-V closely align with established functional gradients, which represent the low-dimensional axes of static functional connectivity across brain regions (Margulies et al., 2016). This spatial consistency between dynamic and static functional organization has been highlighted in previous studies in adults (Abbas et al., 2019; Bolt et al., 2022). Here, we first examined their relationship in developmental cohorts (HCP-D) by comparing the occurrence time of dynamic states and the EVR of static functional gradients. Across all three dynamic states, their occurrence ratio showed strong correlations with the EVR of the corresponding static functional gradients (Fig. 3A, Pattern S-A: *r* = 0.46, *p* < 0.01, Pattern TP-D, *r* = 0.44, *p* < 0.01, Pattern SM-V, *r* = 0.52, *p* < 0.01).

**Figure 3.**
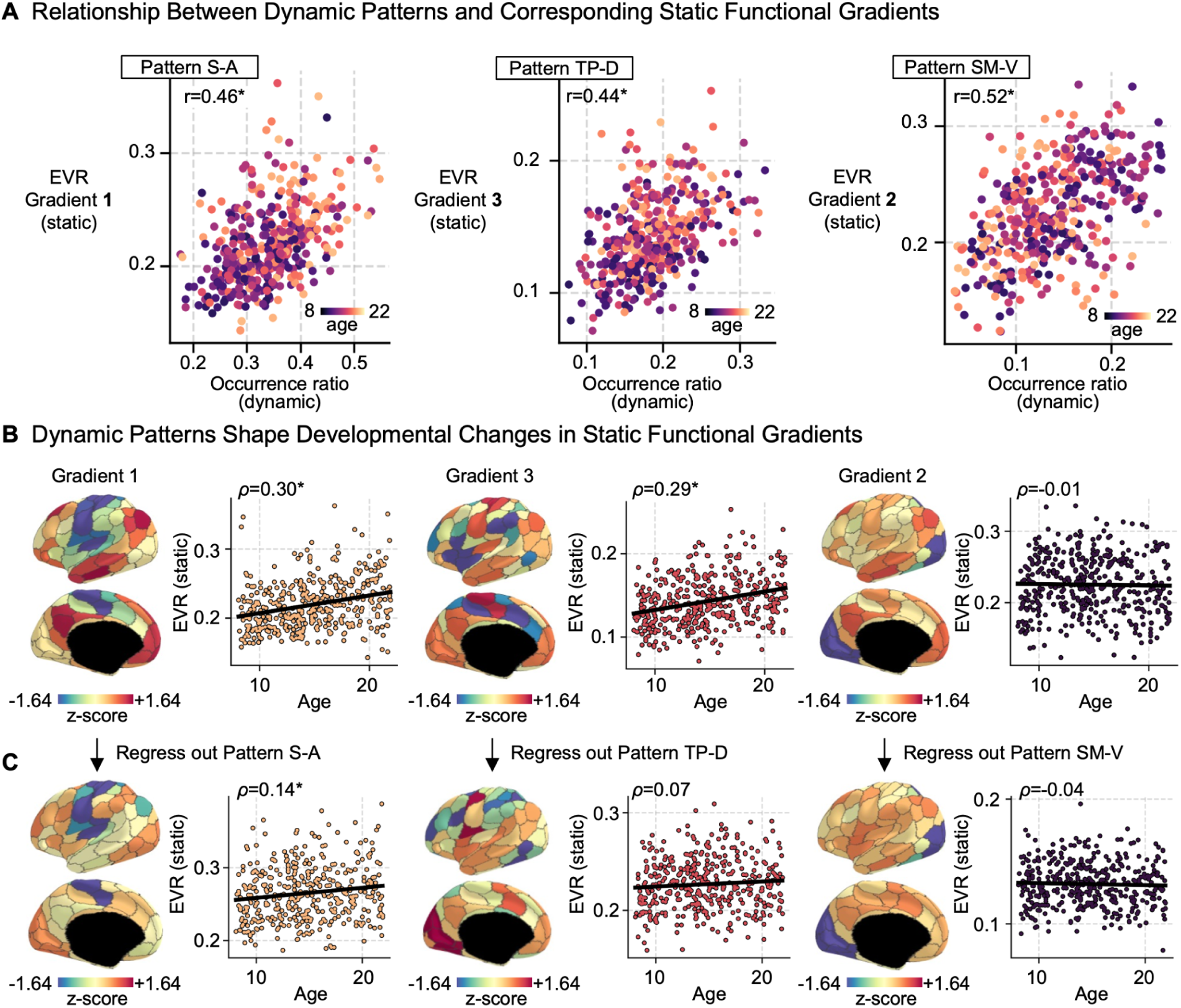
Relationship between the dynamic state and corresponding static functional gradient **(A)** Relationship between the dynamic state and corresponding static functional gradient (x-axis: occurrence ratio of dynamic state, y-axis: explained variance ratio [EVR] of functional gradient). Each dot represents an individual, with age indicated by color. Note that the functional gradient associated with Pattern TP-D is the third most dominant, while Pattern SM-V is the second. **(B-C)** Dynamic patterns contribute to the developmental effect observed in static functional gradients. The brain maps display the functional gradient from the reference cohort (HCP-A). The scatter plot indicates age dependent change in EVR of each gradient. **(B)** The age effect on functional gradient (x-axis: age, y-axis: EVR of the matched functional gradient). **(C)** The age effect decreased after regressing out corresponding dynamic patterns from fMRI data. The brain maps after regressing out propagation components depict the functional gradients derived from the residual rs-fMRI signals. The significance is denoted by asterisks *: *p* < 0.05.

Of note, recent findings demonstrated that static functional gradients undergo significant developmental changes from childhood to adolescence (Dong et al., 2021; Xia et al., 2022). We hypothesize that these developmental changes are supported by the corresponding dynamic states. Specifically, as participants spend more time in a particular dynamic state, that state becomes the dominant gradient in static functional organization. To test this, we examined the age effects on EVR of functional gradients before and after regressing out corresponding dynamic patterns from the fMRI signals (Fig. 3B-C). Notably, regressing out dynamic patterns significantly reduced the age effect on functional gradients, in particular for the functional gradient and pattern along the S-A axis (ρ = 0.30 to ρ = 0.14) (Fig. 3C). These findings provide compelling evidence that the time participants spend in dynamic states plays a critical role in shaping the developmental progression in static functional organization from childhood to early adulthood.

### Increased time spent in top-down propagation and decreased time spent in bottom-up propagation from childhood to early adulthood

The above analyses revealed that the Pattern S-A became more prevalent with age. However, the analysis did not distinguish directionality of propagations that ascend the S-A axis (i.e. bottom-up) and those that descend the S-A axis (i.e. top-down). To address this, we defined ‘bottom-up’ propagation as the temporal changes from sensory to association regions during the 0 to *π* phase and ‘top-down’ propagation as the reverse from association to sensory areas during the *π* to 2*π* phase (Fig. 4A). We then calculated the time participant spent within each propagation phase bin (*π*/16) and assessed its relationship with age. Notably, this analysis revealed an age-related increased prevalence of top-down propagation (i.e. *π* to 2*π*), along with a decrease in time spent in bottom-up propagation, particularly during the later propagation phase (i.e. *π*/2 to *π*) of the bottom-up process as it ascends to the default mode network (Fig. 4B). Additionally, we investigated whether dynamic propagations could predict the total cognitive composite score, a comprehensive measure of crystallized and fluid intelligence (Akshoomoff et al., 2013). Using a PLSR model with 5-fold validation with 1000 iterations, we examined the predictive performance (i.e. r-value between actual and predicted scores) for each of four propagation phase stages. Our results showed that both bottom-up and top-down propagation achieved significant prediction performance (*r_0-_*_*π*_*_/2_* = 0.17 ± 0.04, *p__permutation_* = 0.053; *r*_*π*_*_/2-_*_*π*_ = 0.19 ± 0.03, *p__permutation_* = 0.013; *r*_*π*_*_/2-3_*_*π*_*_/2_* = 0.20 ± 0.03, *p__permutation_* < 0.01; *r_3_*_*π*_*_/2-2_*_*π*_ = 0.24 ± 0.03, *p__permutation_* < 0.01). Given that top-down processing involves goal-directed behavior and selective attention critical for complex cognitive tasks, it appears the top-down propagation (*π* to 2*π* stage) predicts cognitive score better than the bottom-up propagations.

**Figure 4.**
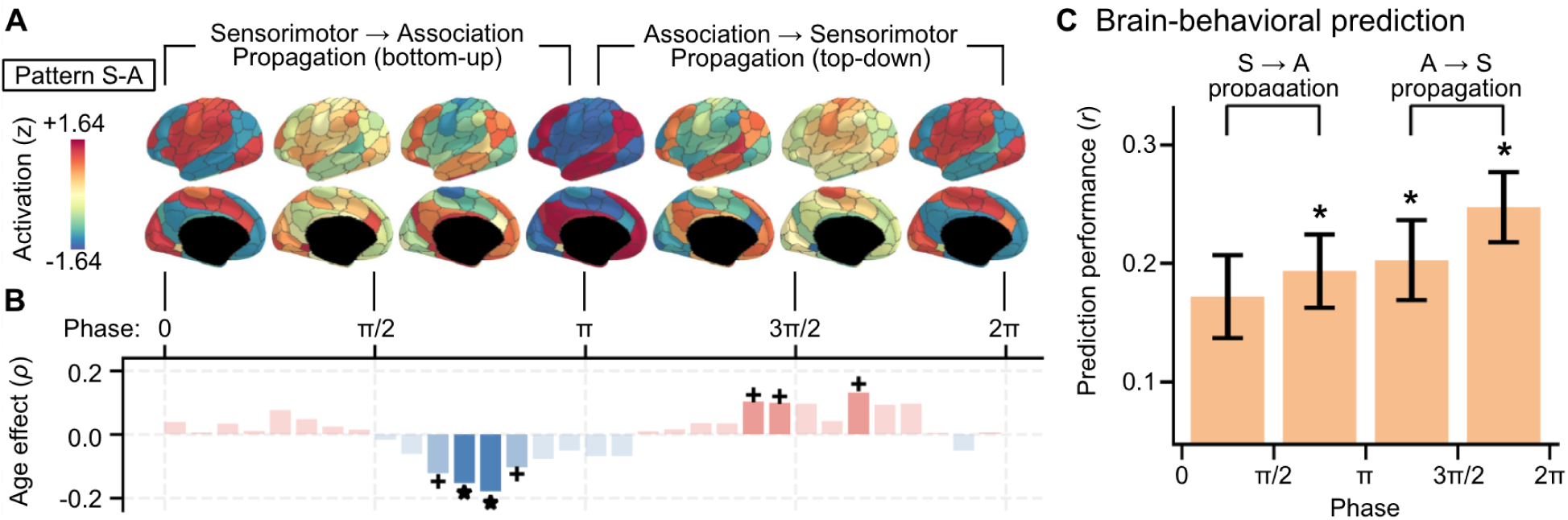
Neurodevelopment of propagation directionality. **(A)** The Pattern S-A derived from the HCP-D cohort, representing group-level propagation patterns. Bottom-up process is defined as the propagation from visual/somatomotor networks to default networks (0 to π) and top-down process (π to 2π) is from default networks to visual/somatomotor networks. **(B)** The age effect of the propagation (y-axis: correlation ρ-value between age and the occurrence ratio at different propagation phases, red bars: positive age effects; blue bars: negative age effects). Statistical significance (+: *puncorrected* < 0.05, \**: pFDR* < 0.05). **(C)** Prediction performance of the composite cognitive score using spatiotemporal patterns for four propagation phase segments (*: *pFDR* < 0.05).

### Developmental shifts in propagation directionality and cognitive relevance of Patterns TP-D and SM-V

Similar to the analysis for Pattern S-A, we also examined age-related changes in propagation directionality for Patterns TP-D and SM-V. For Pattern TP-D, we observed an increasing occurrence of external-to-internal propagation from attention network to default network (*π*/2 to 3*π*/2) and a decreasing occurrence of the reverse internal-to-external propagation from default network to attention network (0 to *π*/2 and 3*π*/2 to 2) from childhood to early adulthood (Fig. S2). For Pattern SM-V, the occurrence increased at the propagation phase from the insular and motor regions to visual networks, while it decreased during the phase from the visual to motor networks. When predicting cognitive scores, both Pattern TP-D and SM-V showed significant but weaker predictive performance compared to Pattern S-A (Fig. S2, Pattern TP-D: r_0-*π*/2_ = 0.21 ± 0.03, r_*π*/2-*π*_ = 0.14 ± 0.03, r_*π*/2-3*π*/2_ = 0.18 ± 0.03, r_3*π*/2-2*π*_ = 0.15 ± 0.03; Pattern SM-V: (r_0-*π*/2_ = 0.15 ± 0.03, r_*π*/2-*π*_ = 0.12 ± 0.04, r_*π*/2-3*π*/2_ = 0.17 ± 0.03, r_3*π*/2-2*π*_ = 0.12 ± 0.04).

### Validation of dynamic propagations and their developmental effects across independent cohorts

The above analyses revealed developmental effects in dynamic propagations from childhood to early adulthood in HCP-D cohort. To evaluate the generalizability of our findings, we replicated our analysis in an independent cohort from the NKI-RS datasets and selected the participants including children, adolescents, and early adults (N = 269, age = 6-21). This dataset consists of three runs with distinct acquisition parameters (TR = 645 ms, 1400 ms, 2500 ms), with the main figures presenting results averaged across acquisition settings for differences in TR. Consistent with our results in HCP-D, the first three propagation patterns explained over 50% variance of resting state fMRI dynamics, each contributing more than 10% of EVR (Fig. S3B). Across cohorts, the propagation patterns demonstrated high similarity compared to the adult reference (HCP-A) (*r* = 0.88 ± 0.04 for Pattern S-A, *r* = 0.81 ± 0.06 for Pattern TP-D, and *r* = 0.87 ± 0.03 for Pattern SM-V). More importantly, showing the similar significant age-related resemblance to adult reference patterns, with Pattern S-A showing the strongest correlation (*ρ* = 0.46, *p* < 0.05) (Fig. 5A, Fig. S3D provides results for each acquisition). Across developmental stages from childhood, adolescent, early adulthood, to middle-age adulthood in NKI-RS, we observed consistent age-related shifts in occurrence ratios in NKI-RS. Specifically, older individuals exhibited a higher proportion of time spent in Patterns S-A and TP-D with a reduced presence in Pattern SM-V (Fig. 5C and Fig. S4 for each acquisition). The correlation between occurrence ratios of dynamic patterns and EVR of static functional gradients also showed high similarities (Fig. S5 for each acquisition). Similarly, we examined age-related changes in occurrence time across different propagation phases in the NKI-RS cohort (Fig. 5D and Fig. S6 for each acquisition). Consistent with HCP-D, a decreased occurrence time was observed in the later phase of bottom-up propagation (*π*/2 to *π*). The increasing trend in the early phase (0 to *π*/2) of bottom-up propagation from HCP-D was more pronounced in NKI-RS, while no significant increase in top-down propagation was detected. This discrepancy might be attributed to the limited data available per individual in the NKI-RS compared to HCP-D. Although we merged occurrence time across three runs to increase data quantity per individual, the NKI-RS dataset have less data per individuals and were collected using varying scan acquisitions, resulting in greater heterogeneity compared to the more standardized HCP-D cohorts. Nevertheless, the overall robustness of the dynamic propagations and their developmental effect was evident across cohorts.

**Figure 5.**
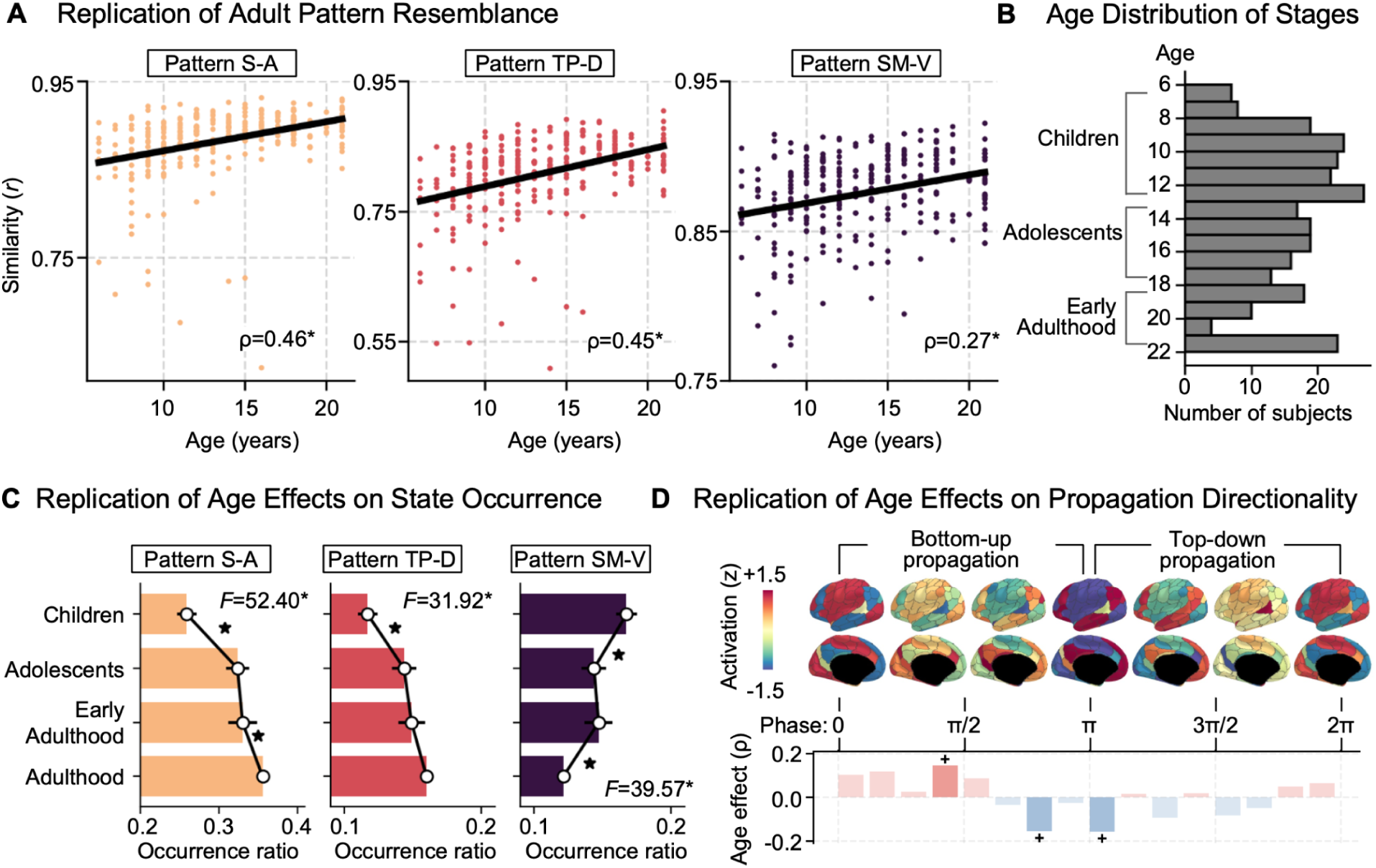
Age effect of dynamic propagation in an independent NKI-RS cohort. **(A)** Age-related similarity of spatiotemporal patterns to the adult reference (HCP-A) across cohorts. **(B)** Age distribution of NKI-RS samples across developmental stages. **(C)** Occurrence ratio of propagation state across developmental stages within NKI-RS. **(D)** Age-related effect (Spearman’s *ρ*-value) on the occurrence ratio at different propagation phases for Pattern S-A (see Fig. S7 for Pattern TP-D and Pattern SM-V).

### Robustness analyses using an alternative parcellation and low-motion subset

To further validate the robustness of our findings, we conducted two replication analyses within the HCP-D cohort: (1) using a different parcellation (Schaefer 400 parcels) (Schaefer et al., 2018), (2) within a low-motion subset consisting of individuals without substantial head movement (N = 287, Mean Framewise Displacement < 0.2 mm. Consistent findings were observed across both replications. Specifically, from childhood to early adulthood, individuals spent more time (i.e. occurrence ratio) for Pattern S-A and Pattern TP-D while less time for pattern SM-V (Fig. S8). The occurrence ratio of the propagation states also showed a strong correlation with the EVR of static functional gradients (Fig. S9). The developmental effect on occurrence ratio at different propagation phases were also replicated across all three propagation patterns (Fig. S10). From childhood to adulthood, participants exhibited a decrease in time spent in bottom-up propagation and an increase in time spent in top-down propagation along the S-A axis. Notably, Pattern S-A showed better predictive performance for the total composite cognitive scores compared to Pattern TP-D and Pattern SM-V, with the highest prediction performance using the top-down propagation in Pattern S-A (Fig. S10). Collectively, these replications highlight the robustness of our results associated with head motion and scale of cortical parcellation.

## Discussion

Mapping the organization of developmental axes is crucial for unraveling how large-scale functional networks mature in the human brain. During early development, major macroscale organizational axes, such as the anterior-posterior gradient, govern cortical neurogenesis, establishing the framework for cortical maturation (Sydnor et al., 2021). As development progresses, the sensorimotor-association (S-A) axis becomes more prominent, shaping intrinsic activity and functional connectome (Dong et al., 2021; Sydnor et al., 2023; Xia et al., 2022). In this study, we build upon these foundational insights by demonstrating the maturation of dynamic propagations along the S-A axis and two additional primary axes. We used a recently developed time-lag dynamic approach, CPCA, to identify three principal axes of brain dynamic propagations: the sensorimotor-association (Pattern S-A), ‘task positive’-default (Pattern TP-D), and somatomotor-visual (Pattern SM-V). Our analyses revealed substantial age-related changes in spatiotemporal patterns, with individuals transitioning from childhood to early adulthood showing increased time spent and higher occurrence rates in Pattern S-A and TP-D dominant states, while spending less time in Pattern SM-V. Additionally, we observed a shift towards top-down propagation, which served as a more accurate predictor of cognitive performance compared to bottom-up propagation. These findings, replicated across independent cohorts with different acquisitions, provide new insights into the evolution of brain dynamics during development and how these changes contribute to cognitive abilities.

Prior studies have consistently identified dynamic patterns of brain function as stable features across differing dynamic analysis approaches – though, primarily in adults (Bolt et al., 2022; Yousefi & Keilholz, 2021). Our study extends these findings to childhood, demonstrating that by age 8, spatiotemporal propagation patterns are already well formed and exhibit high similarity to those observed in adults. Among the three dynamic patterns, Pattern S-A in children most closely resembles the adult pattern and remains the most similar up to adulthood. This pattern follows the hierarchical organization between primary visual/sensorimotor and higher-order association networks, and is associated with electrophysiological changes in infra-slow neural oscillations and high gamma band activity (Buzsáki & Draguhn, 2004; Y. Gu et al., 2021; Majeed et al., 2011). This hierarchical organization also aligns with established trajectories in cortical functional development (Hill et al., 2010). The early emergence of Pattern S-A highlights its critical role in coordinating sensory integration with higher-order cognitive functions, reinforcing its status as a dominant organizational feature for brain dynamics (Dong et al., 2021; Fair et al., 2008; Nguyen et al., 2024; Power et al., 2010; Supekar et al., 2009; Sydnor et al., 2023; Xia et al., 2022). The second propagation, Pattern TP-D, representing interactions between external and internal modes of brain function, also undergoes substantial change during adolescence. These changes reflect the brain’s increasing capacity for internal processing and self-referential thought (Dwyer et al., 2014; Park et al., 2024a). Meanwhile, Pattern SM-V, which propagates between the somatomotor and visual cortices, shows refinement in sensory processing pathways across two primary sensory networks, indicating ongoing maturation and specialization of sensory systems throughout adolescence. Notably, the resemblance to adult dynamic patterns increases linearly from childhood to early adulthood, suggesting a progressively adult-like development of cortical dynamic organization as the brain matures.

One of the most notable findings of our study is the developmental shift in dominant propagation states, with an increase in time spent in Pattern S-A propagation state and a corresponding decrease in Pattern SM-V. This shift reveals how the S-A axis gradually emerges as the primary organizational axis of the intrinsic functional connectome from childhood to adulthood (Dong et al., 2021; Xia et al., 2022). Prior studies have demonstrated a dominant static functional gradient shift from the SM-V axis in childhood to the S-A axis in adolescence, reflecting a reorganization of funcitonal networks (Dong et al., 2021). These functional gradients represent the low-dimension organization of the functional connectivity (Margulies et al., 2016) and their shift parallels developmental changes in behavior. Duuring childhood, the brain predominantly focuses on processing external sensory inputs, while adolescence is characterized by increasing engagement in higher-order cognitive and social activities that require the integration of information pathways across unimodal and multimodal networks, such as theory of mind, memory retrieval, and reasoning (Blakemore, 2008). Our findings expand this understanding from a dynamic perspective, showing that both the S-A and SM-V dynamic propagations are present from early childhood, though the time proportion spent in each dynamic state changes significantly with age. Specifically, participants spent more time in Pattern S-A propagation state and less time in Pattern SM-V propagation state as they progressed from childhood to early adulthood.

This finding has two key implications. First, foundational dynamic propagations along both the S-A and SM-V axes are established early, with their prominence evolving over time. The increasing dominance of the Pattern S-A suggests a gradual shift from a focus on basic sensory processing toward higher-order cognitive integration, paralleling the shift toward more abstract cognitive tasks during adolescence – such as theory of mind, memory retrieval, and reasoning. Second, the increase in occurrence time of Pattern S-A, coupled with a decrease in Pattern SM-V, suggests that developmental shifts in the macroscale organization of static functional connectivity may be driven by changes in dynamic brain activity during youth. The gradual increase in time spent in Pattern S-A may play a key role in establishing the S-A axis as the dominant organizational gradient of the functional network, thereby facilitating the brain’s maturation from childhood to adulthood. Collectively, our finding provides compelling evidence for neurodevelopmental plasticity across timescales. Though, importantly, they also suggest that a substantial portion of age-related differences in static macroscale brain networks observed during development using measures that assume static patterns of functional connectivity, are actually driven by differences in moment-to-moment changes in neural dynamics.

Beyond Pattern S-A, our study further revealed the developmental changes of the propagation in both top-down and bottom-up directions in youth and how this process supports behavioral cognitive function. Directionality of dynamic propagation in the brain can represent the flow of neural activity across different cortical regions, reflecting hierarchical processing where activity moves between higher-order association areas and lower-order sensory regions depending on cognitive demands (Friston, 2005; Mesulam, 1998). The directionality of propagation is critical for cognitive control and executive functions, which undergo significant development during youth to support goal-directed behavior and complex cognitive tasks (Casey et al., 2008; Dalley et al., 2011; Luna et al., 2010). The biological significance of propagation lies in its ability to represent how the brain processes information hierarchically. Previous studies using fMRI have shown that top-down processes are integral to efficient neural function and cognitive development (Y. Gu et al., 2021; Raut et al., 2021; Xu et al., 2023), which become more prevalent as cognitive demands increase as development progresses (Johnson, 2011; Pines et al., 2023). Our findings extend previous work by demonstrating a significant decrease of time spent in bottom-up propagations while an increase in top-down propagations along Pattern S–A was observed as participants matured from childhood to early adulthood, supporting the idea that the brain’s hierarchical processing system becomes more refined and efficient with age (Dosenbach et al., 2010; S. Gu et al., 2015). Importantly, top-down propagation within the S-A axis was found to be a better predictor of cognitive performance compared to bottom-up propagation. However, this prediction was only replicated in datasets with a high quantity of individual scans (totaling > 30 min), emphasizing the importance of sufficient data for measuring individual dynamics. Nevertheless, these findings highlight the role of top-down hierarchical processes in cognitive development, suggesting that the increasing dominance of top-down propagations may reflect the brain’s adaptation to growing cognitive demands. While our results do not directly address neural plasticity or cognitive training, they highlight the importance of understanding how developmental shifts in propagation directionality contribute to cognitive function, which could inform future research on experience-dependent neurodevelopment and targeted interventions.

In addition to Patterns S-A and SM-V, we also identified substantial changes in Pattern TP-D from childhood to adolescence. Dynamic patterns along this TP-D axis reflect the external-to-internal propagations and become increasingly similar to those seen in adults. Similar to Pattern S-A, individuals also spend more time (i.e. increased occurrence ratio) in Pattern TP-D as they age and this higher occurrence ratio of Pattern TP-D is strongly correlated with the static functional organization along the TP-D axis. These findings suggest that, beyond the well-established role of the hierarchical S-A axis, the dynamic propagation along the TP-D axis between frontoparietal/attention networks and default mode network, plays a crucial role in brain development during youth. Previous studies have highlighted the emergence of the TP-D axis as a key axis in thalamocortical connectivity during childhood (Park et al., 2024b). Recent research has also shown that maturational changes in cortical organization are linked to increased within-network connectivity in the ventral attention network, which plays a central role in refining the S-A axis in adolescence (Dong et al., 2024). Additionally, frontoparietal and executive control networks support top-down control and bottom-up feedback processes. During adolescence, these networks undergo notable changes. The control and attention networks transition from interconnection to segregation, a process that supports refining cognitive control abilities (Fair et al., 2009). Task-positive networks were also identified in a unique axis during naturalistic viewing, as compared to the resting state (Samara et al., 2023). This further highlights the role of frontoparietal and attention networks in task-related processing. Throughout adolescence, brain development can be viewed as a process in which brain organization is continuously shaped and refined through various behavioral tasks. The observed increase in external-to-internal propagation ratios may reflect the dynamic, context-dependent reconfiguration of brain networks, enabling a balance between processing external sensory inputs and adapting to the evolving complexity of internal cognitive demands (Dwyer et al., 2014). This reconfiguration is particularly relevant in the maturation of cognitive control, where the interaction between task-oriented networks and the DMN supports the integration of external stimuli with internal cognitive processes to support developmental improvements in executive functioning (Chen et al., 2023). Collectively, these shifting dynamics highlight a fundamental aspect of neural development, underscoring the brain’s transition from reactive to reflective processing.

Notably, the results in our study are robust across various methodological variations and generalizable across an additional independent youth cohort with three different scan acquisitions. At the individual level, CPCA method effectively captured recurring spatiotemporal patterns with high reproducibility and test-retest reliability. Compared to traditional methods such as the sliding-window approach or quasi-periodic pattern analysis with a predefined window size (Majeed et al., 2011), CPCA demonstrates superior precision in detecting the individual dynamic patterns without the need for a fixed arbitrary windowing across individuals from different developmental stages. This makes CPCA particularly flexible and suitable for use with neurodevelopmental studies (Foster & Scheinost, 2024). It’s important to note when studying dynamics, particularly those at the individual level, data quantity remains a critical factor. While most of our findings have been successfully replicated, we did not observe significant results in behavioral prediction for samples with less individual data. Recent fMRI research suggested 20-30 minute data requirements for characterizing individual brain organization, which is even more essential for dynamic studies (Hong et al., 2020; Ooi et al., 2024; Yang et al., 2021). Building on these findings, future research could apply CPCA to naturalistic viewing paradigm, such as movies or music, to examine dynamic brain states in response to continuous, complex, and more ecologically valid stimuli, thereby enriching our understanding of brain interactions in real-world contexts (Eickhoff et al., 2020; Sonkusare et al., 2019; Vanderwal et al., 2019). Expanding the analysis to include subcortical structures and to cover an earlier age range will further enable understanding the emergence of brain organization axes during development (Giedd et al., 1999; Park et al., 2024a; Tau & Peterson, 2010; Xia et al., 2024).

In conclusion, our study revealed the maturation of brain dynamic propagations along key cortical axes from childhood to adolescence. During this critical development stage, time spent in Patterns S-A and TP-D increased, paralleling the shift in functional gradients, suggesting that developmental change of static functional organization may be shaped by moment-to-moment changes in neural dynamics. The increase in top-down propagations along the hierarchical S-A axis emerged as a superior predictor of cognitive abilities, highlighting its role in cognitive maturation. Importantly, the replications across two independent cohorts with different acquisition protocols highlight the robustness and generalizability of our results. Altogether, our study provides compelling evidence that the maturation of brain dynamics support increased cognitive function during adolescence. Future work may build on these findings and examine how functional dynamics adapt in response to moment-to-moment cognitive demands using task-based or naturalistic viewing paradigms. Extending this research to atypical developmental populations and employing longitudinal designs could further offer insights into how brain organization emerges, matures, and may be altered by disease.

## Method

### Datasets

We utilized fMRI data from three large-scale datasets in this study: the Human Connectome Project Development (HCP-D) and Aging (HCP-A), and the Nathan Kline Institute-Rockland Sample (NKI-RS). Our primary analyses were conducted using the HCP-D dataset for the youth sample, with the HCP-A dataset serving as the adult reference. These analyses were then replicated in the NKI-RS dataset. Details of demographic information are summarized in Table S1.

#### HCP-D and HCP-A datasets

are publicly available neuroimaging data from the Lifespan 2.0 Release of HCP project (Bookheimer et al., 2019; Somerville et al., 2018). Resting-state fMRI data were collected across four sites (Harvard, UCLA, University of Minnesota, and Washington University in St. Louis), using 3 Tesla Siemens Prisma scanners. The acquisition protocol is identical for both HCP-D and HCP-A. Specifically, a 2D multi-band gradient-recalled echo (GRE) echo-planar imaging (EPI) sequence was employed (TR = 800 ms, TE = 37 ms, flip angle = 52 degree, with 2 mm isotropic voxel resolution). With this acquisition, during two sessions, two T2*-weighted image runs were acquired, each lasting 6.5 minutes and using different phase encoding directions (AP and PA), resulting in a total of 26 minutes of resting-state fMRI data for each subject. We included the participants whose data had been minimally preprocessed and passed the HCP quality control assessment (Glasser et al., 2013). Our analyses included 408 participants (8-21 years) from HCP-D and a group-average of 399 participants (22-64 years) from HCP-A as adult reference.

#### Nathan Kline Institute-Rockland (NKI-RS)

The NKI-RS dataset (Nooner et al., 2012) is part of a community-ascertained lifespan dataset. Resting-state fMRI data acquired in three acquisitions, each with distinct temporal and spatial resolutions: a 9.7 minute scan with TR = 645 ms at 3-mm isotropic resolution (900 time points, TE = 30 ms, flip angle = 60 degree), a 9-minute scan with TR = 1400 ms at 2-mm isotropic resolution (404 time points, TE = 30 ms, flip angle = 65 degree), and an 5 min scan with TR = 2500 ms at 3-mm isotropic resolution (120 time points, TE = 80 ms, flip angle = 80 degree). The details of the acquisition can be found at https://fcon_1000.projects.nitrc.org/indi/enhanced/mri_protocol.html. After the preprocessing and quality control, individuals aged 6-21 years (N = 298 for TR = 645 ms; N = 288 for TR = 1400 ms; N = 282 for TR = 2500 ms) were included as the youth sample. We also included participants with age 22-64 (N = 526 for TR = 645 ms, N = 533 for TR = 1400 ms, N = 533 for TR = 2500 ms) as an early adult group in replication analysis. A total of 269 young participants (age 6-21 years) and 487 middle-age adults (age 22-64 years) with data from all three runs were included for replication in Fig. 5.

### MRI Data Preprocessing

#### HCP-D and HCP-A datasets

Resting-state fMRI data were preprocessed through the HCP pipeline (Glasser et al., 2013). Briefly, the minimal preprocessing of fMRI data included distortion correction, head movement correction, bias field correction, intensity normalization, registration to a mixture of volume and surface “grayordinate” CIFTI template using multimodal surface mapping (MSMAll), high-pass (> 0.009 Hz) filtering, ICA-FIX denoising, and spatially smoothed with 2-mm FWHM Gaussian kernel. In our study, we further applied global signal regression and temporal filtering to the HCP minimally preprocessed data using the conventional resting-state fMRI low-frequency band-pass range of 0.01-0.08 Hz. Finally, the timeseries were averaged within each parcel using Schaefer 17-network 200 parcellations (Schaefer et al., 2018). To assess the robustness of the findings, Schaefer 17-network 400 parcellation was also employed in our replication analysis (Schaefer et al., 2018). We further replicated the main results using a subset of individuals (N = 287 with low head motion (mean framewise displacement < 0.2 mm) (Power et al., 2014).

#### NKI-RS

Both replication datasets were preprocessed using fMRIPrep (Esteban et al., 2019) and XCP-D pipelines (Mehta et al., 2023). Similar to the HCP pipeline, the functional MRI data pipeline of fMRIPrep and XCP-D includes head motion correction, registration to the “grayordinate” CIFTI template. Nuisance regressors include 36 parameters (six head motion parameters, white matter, cerebrospinal fluid, and global signal, along with their squired terms, derivatives, and squared derivatives) (Ciric et al., 2018). The data were temporally filtered with the same low-frequency range of 0.01-0.08 Hz and parcellated using Schaefer 17-network 200 parcellation (Schaefer et al., 2018).

### Dynamic spatiotemporal pattern analysis

To extract the propagation patterns of resting state fMRI signals, we adopted the CPCA approach (Bolt et al., 2022; Feeny, 2008; Horel, 1984). This method converted the fMRI data into analytic signals followed by a dimensionality reduction technique to decompose the high-dimensional analytic data into low-dimensional principal components. Specifically, for each parcel, we first computed the analytic signal *x_a_(t)* using Hilbert transform as follows:

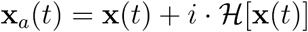

The real part *x(t)* of the analytic signal represents the original fMRI time series, while the imaginary part corresponds to the Hilbert-transformed signal, which is phase-shifted signals by *π/2* radians (Fig. S1A). Next, we applied fast randomized singular value decomposition (SVD: https://github.com/facebookarchive/fbpca) on the complex-valued data matrix X_a_ (size: the number of time points by the number of parcels), decomposing into ten components.

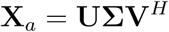

The left singular vectors *U* contain the orthogonal temporal vectors (size: the number of time points by the number of components), Σ is a diagonal matrix which holds the singular values, and the right singular vectors *V* consists of the orthogonal spatial vectors (size: the number of components by the number of parcels) with *H* as Hermitian transpose. These vectors are complex-valued signals which can be represented in terms of amplitude and phase components using Euler’s formula. The amplitude quantifies the strength of the components, and the phase specifies the relative delay, expressed in radians from 0 to *2π*. To capture a full cycle of cortical propagations, we extracted the spatiotemporal patterns across 32 equally spaced phase segments (i.e. 2*π/32*), yielding 32 spatial maps at each phase segment from 0 to 2*π* for each component. The selection of 32 phase bins achieves a balance between computational efficiency and temporal resolution in the phase domain. Specifically, we first averaged the temporal vector U for each component within each of 32 phase segments to generate the resampled temporal vector (32 by the number of components). The spatiotemporal patterns were then computed by performing a component-wise outer product between the resampled temporal vector and the corresponding spatial vector. In Fig. 1A, we demonstrated and visualized spatial maps at phase 0, 5*π*/32, 10*π*/32, 15*π*/32, 20*π*/32, 25*π*/32, and 30*π*/32 for the first three components.

To improve inter-individual comparability, we first generated the adult components from the HCP-A cohort to serve as a reference for alignment. We concatenated fMRI data across individuals in HCP-A cohort and performed the SVD analysis on the analytic fMRI signal to extract the temporal and spatial vectors as adult reference. We then used the Procrustes matching to align spatial vectors (size: the number of parcels by the number of components) between each individual from the HCP-D cohort and the reference (Schönemann, 1966; Schultz et al., 2014). The alignment transformation (rotation and reflection) was subsequently applied to the temporal vectors to match with the aligned spatial vectors for each individual. Adult resemblance was quantified using a Pearon’s correlation between each individual’s flattened propagation pattern and the group-reference adult pattern.

### Occurrence ratio of dynamic state and age effect analysis

To assess the time participants spent (i.e. occurrence ratio) in each of the propagation states defined by the CPCA components, each time point of the cortical fMRI frames was assigned to one of the CPCA components. The temporal strength of the components at every time point was quantified using weighted temporal vector (i.e. *U*Σ, size: the number of time points by the number of components). The moment-by-moment propagation state was determined using a ‘winner-takes-all’ strategy, where the state with the highest temporal magnitude (Fig. 2A) at each time point was assigned as the dominant state. Here, temporal magnitude was calculated as the amplitude of the temporal vector weighted by the singular value of the corresponding propagation pattern at that moment. For the time points where none of the first three components were dominant, we grouped them as “other patterns” state. For each individual, the occurrence ratio was calculated as the proportion of time points during which a specific state was dominant, relative to the total number of time points of fMRI scans.

To compare the occurrence ratio across the developmental stages from childhood to adulthood, participants from the HCP-D were first grouped into three developmental stages within dataset children (8 ≤ age < 13), early adolescents (13 ≤ age < 18), and late adolescents (18 ≤ age < 22). The late adulthood stage was measured using HCP-A dataset. ANOVA was used to compare the difference across four age groups, followed by sequential pairwise comparisons between age stages using t-tests. To account for multiple comparisons, we applied the Benjamini-Hochberg procedure to control the false discovery rate in the Python package ‘statsmodels’ (Seabold & Perktold, 2010).

### Dynamic directionality analysis

To assess the age effect on the directionality of dynamic propagations along each CPCA component, we counted the time (i.e. occurrence ratio) that participants spent in each of the 32 phase bins (2*π/32*). Next, we calculated the Spearman’s correlation between age and occurrence ratio for each phase bin. For Pattern S-A, propagations during the 0 to *π* phase captured spatial transitions from visual and somatomotor to default mode regions (i.e. bottom-up), while the *π* to 2*π* phase captured a reversed, top-down propagation. We applied the same approach to assess the age effect on occurrence ratios within each propagation phase bin for Patterns TP-D and SM-V.

### Brain-wide prediction

In this study, the brain-wide prediction was conducted using partial least squares regression (PLSR) with stratified five-fold cross-validation to ensure a balanced distribution of the dependent variable (Krishnan et al., 2011; McIntosh & Mišić, 2013). In PLSR, we z-scored the input features for normalization and set the hyperparameter *n_components*=3 in the *‘scikit-learn’* Python package (Pedregosa et al., 2011). To examine the extent to which age can be represented by all three dynamic patterns in youth, we flattened spatiotemporal patterns of three components (32 spatial maps by 3 components) and concatenated them as input features. Model performance was evaluated by calculating the mean square error and Spearman’s correlation between actual and predicted age in the HCP-D dataset.

A similar approach was used for brain-behavior prediction of cognitive scores. To identify which propagation stage (phase segment) best predicts cognitive performance, weperformed separate predictions for each component and each propagation stages (0 to *π*/2, *π*/2 to *π*, *π* to 3*π*/2, and 3*π*/2 to 2*π*), also using PLSR with five-fold cross-validation. The prediction was repeated 1,000 times to ensure robustness of the prediction score. To asssess statistical significance, a null-distribution was generated using a permutation test (1,000 iterations) by randomly shuffling cognitive scores. The mean observed prediction performance was then compared against this null distribution to determine statistical significance.(Pedregosa et al., 2011).

### Functional gradients of functional connectivity

To obtain the low-dimensional representation of the static functional connectome, we calculated the first ten functional gradients of the functional connectivity for each individual. Previous studies have demonstrated that gradients derived from the principal component analysis (PCA) provided higher reliability and prediction power than the traditional diffusion embedding method (Hong et al., 2020). Accordingly, we used PCA to extract the functional gradients here. Specifically, we computed the adjacency matrix by applying Pearson’s correlation to the row-wise thresholded (top 10%) functional connectivity matrix using the *BrainSpace toolbox* (Vos de Wael et al., 2020). Procrustes alignment (rotation and reflection) was then applied to match the individual gradients with the reference gradient derived from the averaged functional connectivity matrix of the HCP-A cohort. Finally, we compared the EVR of the functional gradients with that of the propagation components using Spearman’s correlation.

### Reliability analysis

To examine the test-retest reliability of the spatiotemporal patterns, we employed a multivariate non-parametric method, discriminability (Bridgeford et al., 2021). For the HCP-D cohort, each subject had four rs-fMRI scans with two different phase encoding directions, we first concatenated two scans with the same phase encoding direction, and then evaluated the reliability between two resulting concatenated subsets. For the NKI-RS cohorts, the test-retest reliability analysis was assessed by comparing pairs of group-level spatiotemporal patterns from rs-fMRI scans acquired at TR = 645 ms, TR = 1400 ms, and TR = 2500 ms.

### Replication analysis in NKI-RS

To evaluate the generalizability of our findings, we replicated the above analyses, including dynamic spatiotemporal pattern analysis, occurrence ratio, and directionality analysis using the NKI-RS dataset. The dataset were collected across three runs during the same session, each with different acquisition parameters (TR = 645 ms, 3 mm isotropic, 9.7 min; TR = 1400 ms, 2 mm isotropic, 9.4 min; TR = 2500 ms, 3 mm isotropic, 5.0 min). We calculated the spatiotemporal patterns for each run separately and replicated the findings in Figure S4-7. To increase data quantity robustness estimation per individual, we also merged three runs together by averaging spatiotemporal patterns and occurrence ratio weighted by the scan duration per run for each individual.

## Data and code availability

The Human Connectome Project datasets are publicly available from the Lifespan HCP Release 2.0 (available at https://www.humanconnectome.org/study/hcp-lifespan-aging/article/lifespan-20-release-hcp-aging-hcp-development-data) hosted on the NIMH Data Archive (NDA) (https://nda.nih.gov/). Data from the NKI-RS study are available through the Database of the International Neuroimaging Data-sharing Initiative (INDI) (https://fcon_1000.projects.nitrc.org/indi/enhanced/). The analyses in this study utilized publicly available cortical atlases, specifically the Schaefer-200 and Schaefer-400 atlases, which can be accessed at https://github.com/ThomasYeoLab/CBIG/tree/master/stable_projects/brain_parcellation/Schaefer2018_LocalGlobal.

The preprocessed data for NKI-RS dataset was available from the Reproducible Brain Charts (RBC) project through https://reprobrainchart.github.io/. All subsequent analyses and statistical computation were carried out in Python 3.11.9 (https://www.python.org/). The complex principal component analysis used in this study was adopted from https://github.com/tsb46/complex_pca. The code for the analyses in our study is provided at https://github.com/HumanBrainED/Neurodev-CPCA.

## Acknowledgements

This work was supported by grants from the National Institute of Health (NIH) awards R01MH139349, RF1MH128696, and P50MH109429 to T.X. K.B. was supported by Basic Science Research Program through the National Research Foundation of Korea (NRF) funded by the Ministry of Education (RS-2024-00407857). This work is also supported by gifts from Joseph P. Healey, Phyllis Green, and Randolph Cowen to the Child Mind Institute and the National Institutes of Health fundings (R24MH114806 to M.P.M.). T.D.S. was supported by the NIH (R01MH113550, R01MH112847, R37MH125829, R01MH120482, R01EB022573), the Penn-CHOP Lifespan Brain Institute, and the AE Foundation. H.P. was supported by National Research Foundation of Korea (RS-2024-00408040), Institute for Basic Science (IBS-R015-D2), Artificial Intelligence Innovation Hub program (RS-2021-II212068), ICT Creative Consilience program (IITP-2025-RS-2020-II201821), and AI Graduate School Support Program (RS-2019-II190421). S.J.H. was supported by the Institute for Basic Science IBS-R015-D1, the National Research Foundation of Korea (NRF) grant funded by the Korea Government (MSIT) (NRF-2022R1C1C1007095, RS-2023-00217361, RS-2024-00398768 to S.J.H.)

## Notes

### Competing Interest Statement

The authors have declared no competing interest.

